# Temporal inhibition of chromatin looping and enhancer accessibility during neuronal remodeling

**DOI:** 10.1101/2021.08.30.458233

**Authors:** Dahong Chen, Catherine E. McManus, Behram Radmanesh, Leah H. Matzat, Elissa P. Lei

**Affiliations:** Nuclear Organization and Gene Expression Section, Bethesda, MD, USA; Laboratory of Biochemistry and Genetics, National Institute of Diabetes and Digestive and Kidney Diseases, National Institutes of Health, 9000 Rockville Pike, Bethesda, MD, 20892, USA; Laboratory of Cellular and Developmental Biology, National Institute of Diabetes and Digestive and Kidney Diseases, National Institutes of Health, 9000 Rockville Pike, Bethesda, MD, USA

**Keywords:** enhancer-promoter looping, chromatin accessibility, neuronal remodeling, Shep, anti-looping

## Abstract

During development, looping of an enhancer to a promoter is frequently observed in conjunction with temporal and tissue-specific transcriptional activation. The chromatin insulator-associated protein Shep promotes *Drosophila* post-mitotic neuronal remodeling by repressing transcription of master developmental regulators, such as *brain tumor* (*brat*), specifically in maturing neurons. Since insulator proteins can promote looping, we hypothesized that Shep antagonizes *brat* promoter interaction with an as yet unidentified enhancer. Using chromatin conformation capture and reporter assays, we identified two novel enhancer regions that increase in looping frequency with the *brat* promoter specifically in pupal brains after Shep depletion. The *brat* promoters and enhancers function independently of Shep, ruling out direct repression of these elements. Moreover, ATAC-seq in isolated neurons demonstrated that Shep restricts chromatin accessibility of a key *brat* enhancer as well as other enhancers genome-wide in remodeling pupal but not larval neurons. These enhancers are enriched for chromatin targets of Shep and are located at Shep-inhibited genes, suggesting direct Shep inhibition of enhancer accessibility and gene expression during neuronal remodeling. Our results provide evidence for temporal regulation of chromatin looping and enhancer accessibility during neuronal maturation.

## INTRODUCTION

Establishment and maintenance of proper chromatin topology has emerged as an essential and conserved feature of exquisitely tuned, developmentally regulated gene expression programs. On the finest scale, looping between an enhancer and a promoter (E-P looping) is frequently observed during or even before the onset of developmentally programmed transcriptional activation^1,2^. In fact, forced E-P looping can result in ectopic activation of gene expression^3,4^, suggesting that the E-P looping step itself may serve as a point of either positive or negative regulation during development. One such E-P loop-promoting factor is the Lim domain-containing protein Ldb1, which is expressed during the development of specific tissues and forms a complex with particular transcription factors^5^. Although perturbation of transcription factors may indirectly affect E-P loop formation by affecting transcription^6-8^, no such dedicated antagonist of E-P looping has yet been identified. Furthermore, regulation of chromatin 3D structure during post-mitotic neuronal remodeling has not previously been studied.

Architectural proteins, such as insulator proteins, have been demonstrated to participate in the formation of topologically associating domains and cell type-specific E-P loops. The first tissue-specific regulator of insulator activity to be identified is the *Drosophila* RNA-binding protein Alan Shepard (Shep), which acts as an insulator antagonist only in the nervous system^9,10^. Shep is required for neuronal remodeling, an essential and conserved process that replaces juvenile neuronal projections with adult-specific projections during the metamorphic transition between larval and pupal development. Shep functions in part by repressing transcription of a key neuronal remodeling inhibitor, *brat*, specifically in pupal neurons^11-13^. Shep associates with the chromatin of *brat* and many other target genes^11^, frequently at promoters^10^. We therefore speculated that Shep may antagonize *brat* E-P looping in order to repress its transcription during post-mitotic neuronal remodeling.

## RESULTS

### Shep inhibits *brat* promoter looping with proximal genomic regions

In order to survey Shep-dependent *brat* promoter looping, we performed Circularized Chromosome Conformation Capture (4C-seq) in central nervous system-derived BG3 cultured cells. Shep inhibits transcription of all *brat* isoforms in BG3 cells and pupal neurons (Fig. 1A-C), and ChIP-seq of Shep indicates chromatin association within the immediate vicinity of each of the annotated *brat* promoters^11^ (Fig. 1A). Since *brat-F* is the dominant isoform in pupal neurons^11^ (Fig. 1A), a validated Shep chromatin binding site at the *brat-F* promoter (Fig. S1) was selected as an anchor to generate 4C-seq libraries (Fig. 1A). Upon efficient knockdown of Shep (Fig. 1D), we identified two regions within 30 kb of the anchor that display statistically increased interaction with the *brat-F* promoter (Fig. 1A and E, regions 1 and 3) as well as a third region with decreased interaction frequency (region 2). We hypothesized that these regions could harbor *cis*-regulatory elements, such as enhancers, that loop to the *brat-F* promoter to activate its expression. Furthermore, we speculate that Shep may antagonize these putative E-P looping interactions in a temporally-regulated manner.

**Fig. 1.**
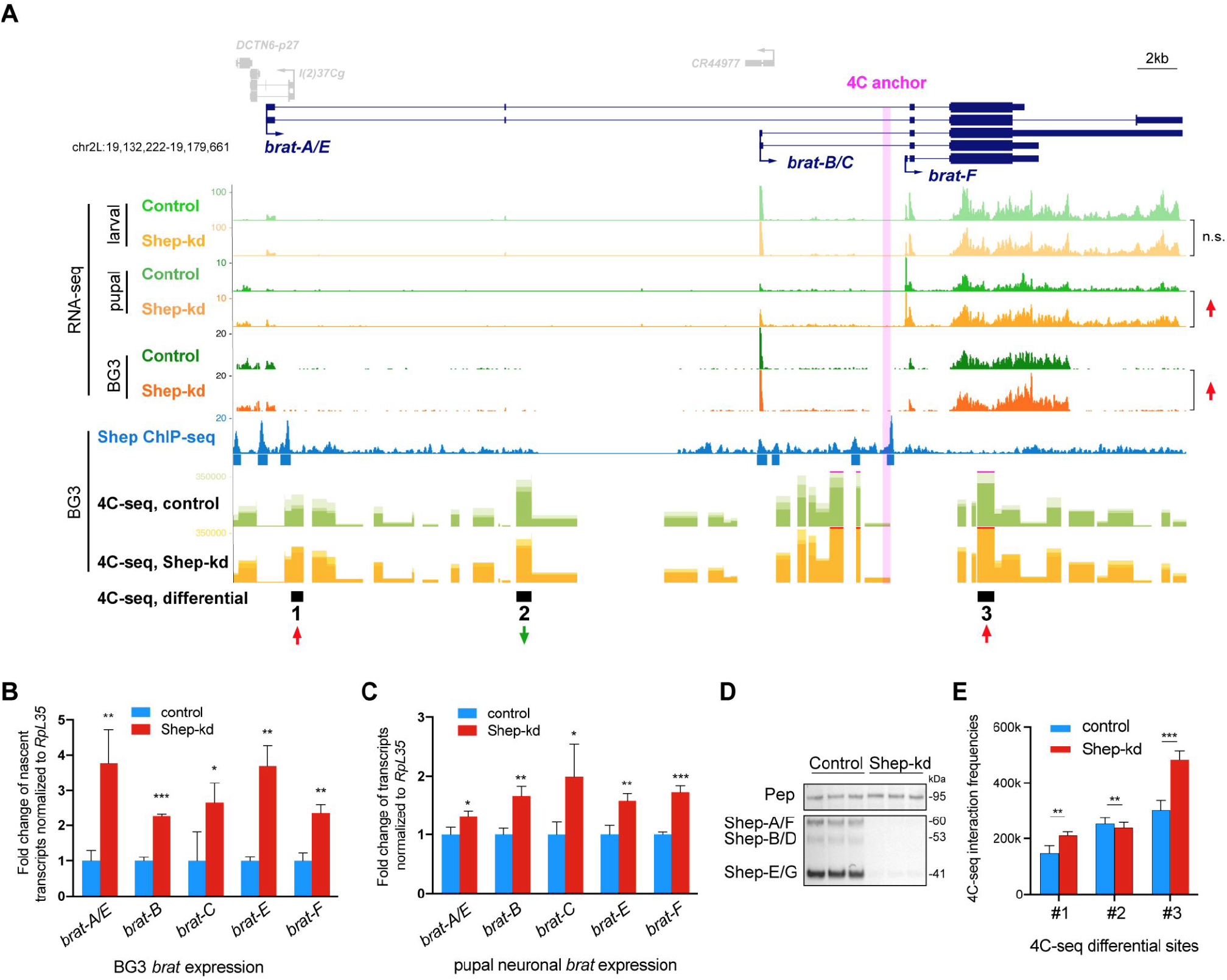
Shep inhibits *brat* expression and *brat-F* promoter looping with proximal genomic regions. **(A)** RNA-seq analysis of control versus Shep-depleted larval neurons, pupal neurons (FDR=1.6e-3, fold change=1.4), and BG3 cells (FDR=5.6e-2, fold change=1.2) at the *brat* locus (top). Note different scales of RNA-seq tracks between larval and pupal neurons, indicating a dramatic decrease in *brat* expression during neuronal remodeling. ChIP-seq profile of Shep in BG3 cells with called peaks indicated below (middle). BG3 cell 4C-seqs using the *brat-F* promoter as an anchor (pink shading) identifies three regions (red and green arrows) with differential (FDR=7.0e-3, 5.6e-3 and 1.4e-4, respectively; fold change=1.3, 0.8 and 1.3, respectively) interaction frequencies upon Shep depletion (bottom). Note that only these three regions pass the FDR < 0.01 threshold. **(B)** Shep depletion in BG3 cells leads to increased transcription of all *brat* isoforms. Nascent RNA was quantified by EU-qPCR using isoform-specific primers. **(C)** Shep depletion leads to increased steady-state expression of all *brat* isoforms in sorted pupal neurons. **(D)** Efficient Shep protein depletion achieved by dsRNA treatment in BG3 cells. Pep served as a loading control for Western blotting. **(E)** Graph showing actual looping frequency measured by 4C-seq in panel A is displayed specifically for the three regions displaying differential interaction that pass the statistical significance threshold.

### Shep inhibits *brat* promoter looping to a neural enhancer

We observed that all three differentially interacting 4C-seq regions are covered by histone post-translational modifications associated with enhancer activity. Examination of publicly available H3K4me1, H3K27ac, and STARR-seq profiles in BG3 cells^14,15^ provided evidence that these regions may function as enhancers to regulate *brat* transcription (Fig. 2A). In order to test this possibility, we cloned each of the three putative enhancer regions juxtaposed to the *brat-F* promoter upstream of a firefly luciferase reporter (Fig. 2B) to assay luciferase activity driven by these constructs or by the *brat-F* promoter alone. We found that only region 1 preceding the *brat-F* promoter is able to increase reporter expression in BG3 relative to control constructs (Fig. 2C), whereas region 1 alone does not support substantial luciferase expression (Fig. S2A-B). Interestingly, none of the constructs are able to enhance *brat-F*-dependent expression when transfected into S2 cells, a non-neural cell type (Fig. 2D and S2C), indicating that region 1 is a cell type-specific enhancer.

**Fig. 2.**
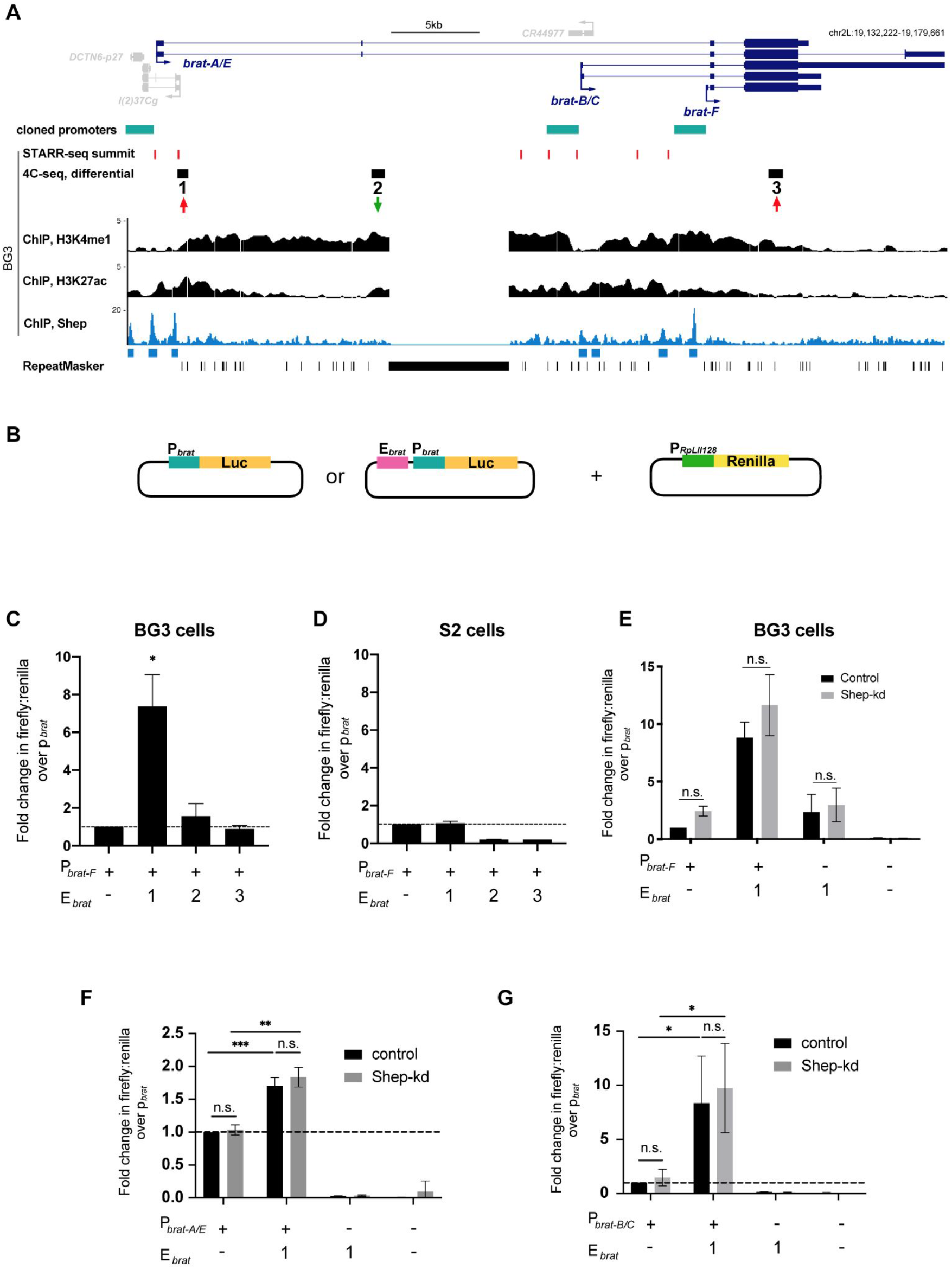
Region 1 is a neural-specific enhancer, and Shep does not directly repress *brat* enhancer or promoter activities. **(A)** ChIP-seq of H3K4me1, H3K27ac, and Shep along with STARR-seq summits at the three 4C-seq differential regions in BG3 cells (red and green arrows). The *brat* promoters are defined as the 2 kb region upstream of the transcription start site of *brat* isoforms and are indicated by teal bars. **(B)** Luciferase constructs with *brat-F* promoter and/or putative enhancer region co-transfected along with a Renilla construct with *RpLII128* promoter, which serves as a transfection control for enhancer reporter assays. **(C-D)** Renilla-normalized luciferase expression driven by the *brat-F* promoter in BG3 or S2 cells. Region 1, but not regions 2 and 3, enhances luciferase expression in BG3 but not S2 cells. **(E)** Normalized luciferase expression in control or Shep-depleted BG3 cells. Expression of region 1 enhancer and/or *brat-F* promoter constructs are unaffected by Shep depletion. Student’s *t* test. **(F)** Fold change of Renilla-normalized luciferase expression over *brat-A/E* promoter driven by region 1 or *brat-A/E* promoter alone or region 1 alone in BG3 cells. Cells were co-transfected with GFP (control) or *shep* dsRNA. **(G)** Same assay as (F) for *brat-B/C* promoter. Student’s *t* test, \**p*<0.05, \*\**p*<0.01, \*\*\**p*<0.001. Average values are reported as mean±sd for all luciferase assays.

We next tested whether Shep affects luciferase expression in this artificial context, in which the region 1 enhancer is juxtaposed to the *brat-F* promoter. We repeated the reporter assays with the *brat-F* promoter alone, juxtaposed to the region 1 enhancer, or the region 1 enhancer alone in control versus Shep-depleted cells. Importantly, no significant differences were observed in control versus Shep-depleted cells for any of these constructs, indicating that Shep does not directly affect either *brat-F* promoter or region 1 enhancer activities (Fig. 2E). Since Shep associates with regions at or nearby promoters of other *brat* isoforms as well, we also performed reporter assays to test Shep regulation of *brat-*A/E or *brat-*B/C promoters with or without region 1. While region 1 can activate both promoters, depletion of Shep does not affect activity of either promoter, indicating that each of these elements also functions independently of Shep in this artificial context (Fig. 2F and G). These key results suggest that Shep does not simply act as a transcriptional repressor of either enhancer or promoter activities. Taken together, we conclude that Shep repression of *brat* only occurs in the *in vivo* context, in which we hypothesize that Shep-mediated antagonism of E-P looping in pupae attenuates *brat-F* expression.

### Shep inhibits neural *brat* promoter-enhancer looping in a stage-specific manner

In order to further test this hypothesis, we next asked whether Shep inhibits *brat-F* promoter looping with the region 1 enhancer *in vivo* in a stage-specific manner. We performed directed 3C using Taqman qPCR to quantify looping between the *brat-F* promoter anchor and the surrounding vicinity, including regions 1-3, in isolated control pupal brains versus brains harboring neurons depleted of Shep (*elav>Dcr-2, shep-RNAi, mCD8::GFP*) (Fig. 3A). Consistent with our 4C-seq results in BG3 cells (Fig. 1A), we found increased looping in Shep-depleted pupal brains between the *brat-F* promoter and the region 1 enhancer as well as a flanking region, which we named region 4 (Fig. 3A). In contrast, larval brains show no statistically significant differences for *brat-F* promoter interaction frequencies with regions 1 and 4, with low looping interactions in both control and Shep-depleted flies. Interaction frequencies in both larvae and pupae are also low between the *brat-F* promoter and other sites examined, including regions 2 and 3. We further interrogated looping in pupal brains using the *brat*-*B/C* promoter as an anchor but found very low levels of looping with regions 1 and 4 relative to looping observed with the *brat-F* promoter (Fig. S3), demonstrating a correlation between low looping frequency and low expression of *brat-B/C*. Moreover, Shep depletion in pupal brains has no effect on looping of regions 1 and 4 with the *brat-B/C* promoter (Fig. S3). Examination of Shep chromatin association in the larval brain revealed that Shep does associate with chromatin at the *brat-F* promoter as well as regions 1 and 4 (Fig. 3A), suggesting direct function at these sites. Given that Shep inhibits *brat* expression in neurons specifically at the pupal stage, our results provide strong evidence for Shep inhibition specifically of *brat-F* looping with enhancer regions 1 and 4 in a temporal manner.

**Fig. 3.**
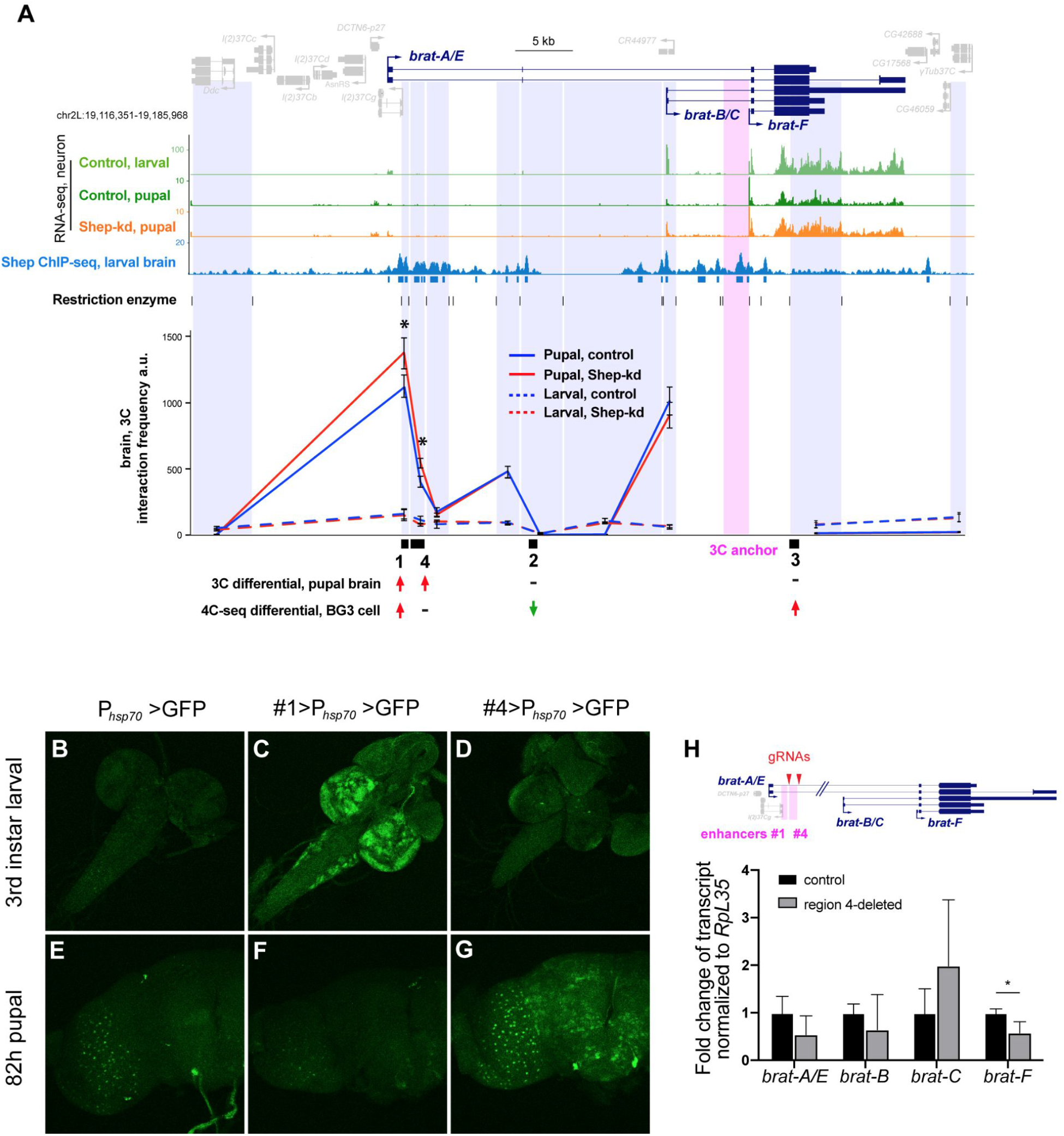
Shep inhibits *brat* E-P looping in a stage-specific manner. **(A)** 3C interaction frequencies in control (blue) or Shep-depleted (red), larval (dashed) or pupal (solid) brains using the *brat-F* promoter as an anchor (pink shading). Digested genomic fragments for 3C quantification are labeled in purple shading. Normalized interaction frequencies in arbitrary units are analyzed by Student’s *t* test. Statistical significance is presented as \**p<*0.05 for three biological replicates. **(B-G)** Regions 1 and 4 harbor temporally-regulated enhancer activities. GFP reporter expression driven by the minimal *hsp70* promoter with or without candidate enhancer regions was visualized by immunostaining larval and pupal brains with anti-GFP. Note leaky GFP expression driven by *hsp70* promoter in pupal brains (E). All flies were grown at 25°C. **(H)** Quantification of steady state *brat* expression in pupal brains of homozygous CRISPR-deletion of region #4 mutants by RT-qPCR. The gRNAs used to target region 4 labeled with arrow heads.

### Region 4 functions as a pupal enhancer *in vivo*

We thus suspected that both regions 1 and 4 may act as enhancers *in vivo*, so we cloned regions 1 and 4 individually juxtaposed to the *hsp70* promoter upstream of a GFP reporter to test their ability to activate transcription in flies. We inserted these constructs into the *attP40* docking site using *PhiC31* integrase in order to assay GFP expression driven by regions 1 or 4 compared to the *hsp70* promoter alone. We observed that region 4 strongly activates GFP expression specifically in pupal but not larval brains (Fig. 3D and G). To test its *in vivo* activity, we CRISPR-deleted the region 4 enhancer in flies and observed reduction of *brat-F* expression in pupal brains (Fig. 3H), indicating a critical function for region 4 as an enhancer of *brat* transcription at this stage. On the other hand, region 1 enhances *hsp70* promoter-driven GFP expression only in larval but not pupal brains (Fig. 3C and F). Consistent with the luciferase reporter assays in BG3 cells, enhancer activities of regions 1 and 4 were unchanged in strong loss-of-function Shep mutant brains (Fig. S4A-D), indicating that Shep does not simply repress enhancer activities. In conclusion, both regions 1 and 4 function as *brat* enhancers *in vivo*, in larvae and pupae respectively, and Shep cannot directly repress the activity of either enhancer. These results are consistent with the hypothesis that Shep functions as an antagonist of E-P looping in order to negatively regulate *brat* transcription at the pupal stage.

### Shep inhibits chromatin accessibility of the region 4 enhancer in pupal neurons

In order to gain insight into the mechanism by which *brat* E-P looping is regulated by Shep, we performed ATAC-seq to examine chromatin accessibility in control versus Shep-depleted neurons. To this end, neurons from control larvae or pupae were GFP-labeled (*elav>Dcr-2, mCD8::GFP*) and FACS-isolated for Omni-ATAC-seq analyses^16^. Consistent with overall decreased expression between larval and pupal stages in control neurons, chromatin accessibility of *brat* overall is also strongly reduced during this developmental transition (Fig. 4A). In contrast, chromatin accessibility increases at the *brat-F* promoter in pupae, perhaps related to primary use of this promoter at the pupal stage. Importantly, in pupal but not larval neurons (Fig. 4A and Table S1), Shep depletion leads to elevated accessibility both in the vicinity of the *brat-F* promoter and at the region 4 enhancer. These results indicate close correspondence between Shep inhibition of chromatin accessibility and Shep inhibition of *brat* E-P looping and transcription in pupae.

**Fig. 4.**
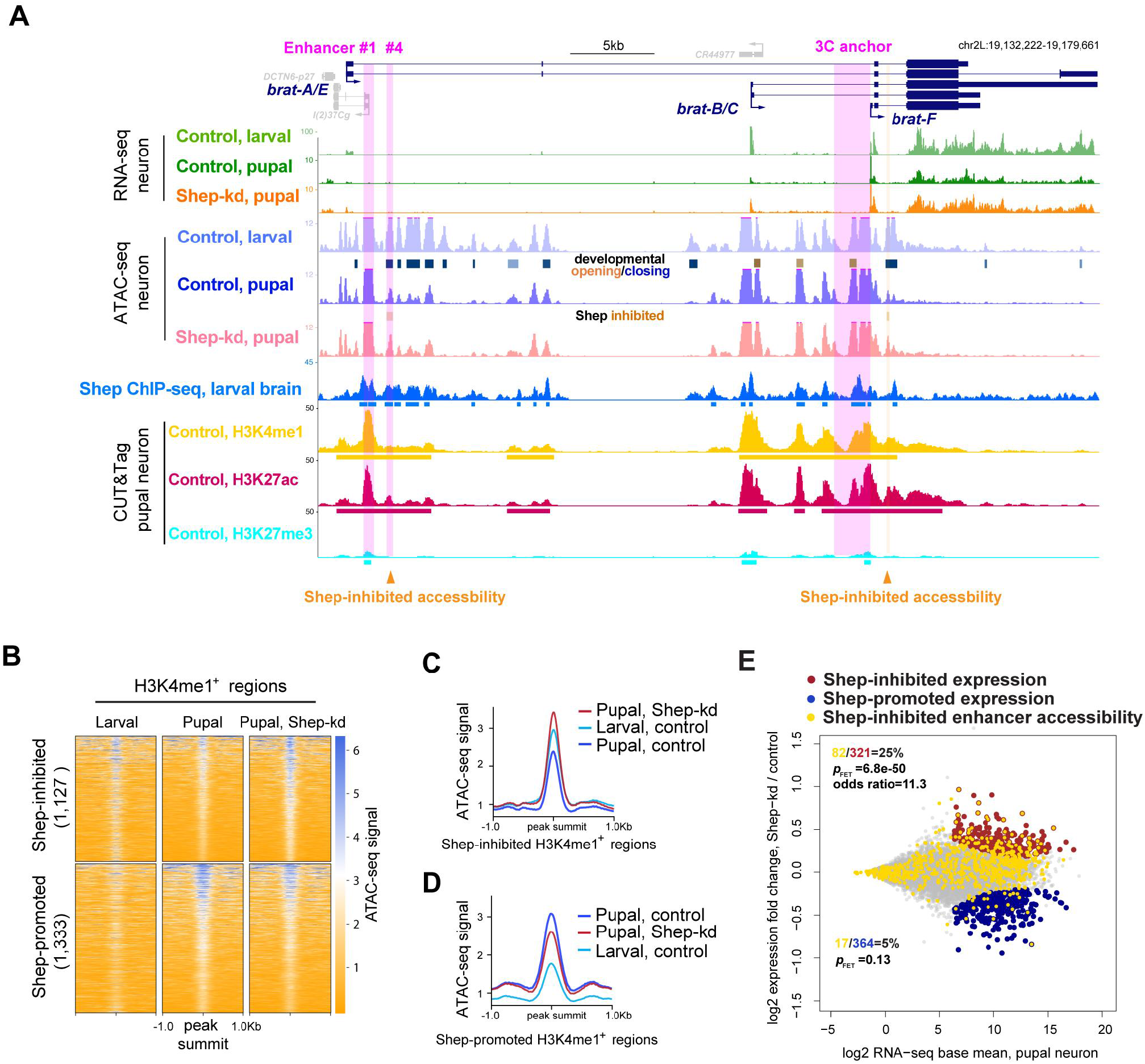
Shep mainly represses enhancer accessibility in pupal neurons genome-wide. **(A)** ATAC-seq profiles of control larval, control pupal, and Shep-depleted pupal neurons at the *brat* locus. Called differentially accessible regions that open (orange) or close (blue) between control larvae and pupae are indicated. Only two accessible regions change (open) after Shep depletion in pupal neurons (orange arrow heads and shading; FDR=0.07 and fold change=1.3 for both regions). CUT&Tag signals of H3K4me1, H3K27ac, and H3K27me3 in sorted control pupal neurons are shown in yellow, red, and cyan. **(B)** ATAC-seq signals are shown for Shep-regulated ATAC-seq peak regions in control versus Shep-depleted pupal neurons that are also marked by CUT&Tag signal for H3K4me1 in sorted control pupal neurons. Corresponding signals in control larval neurons are also shown. **(C-D)** Average ATAC-seq signals across regions in B are shown. **(E)** Shep inhibits enhancer accessibility to repress gene expression in sorted pupal neurons. Genes corresponding to Shep-regulated enhancer accessibility are labeled in gold and overlaid with genes for which expression is Shep-inhibited in red or Shep-promoted in blue. Numbers of genes in each group are indicated and colored accordingly, and FET is reported for indicated colored groups.

### Shep facilitates developmental closing of putative enhancers

Our ATAC-seq profiles enabled us to identify developmentally programmed changes in chromatin accessibility of larval versus pupal control neurons genome-wide, beyond the *brat* locus. The majority of ATAC-seq peaks (33,882 of 41,568) display differential accessibility between control larval and pupal neurons, resulting in 17,367 opening and 16,515 closing regions over this time window. In order to determine whether accessibility changes occur at enhancers genome-wide during neuronal remodeling, we performed CUT&Tag of H3K4me1 on FACS-sorted control larval and pupal neurons (*elav>Dcr-2, mCD8::GFP*) to identify all putative enhancers irrespective of activity. Intersection between ATAC-seq and CUT&Tag profiles indicates that 70% (24,155 out of 33,882) of temporal accessibility changes occur at H3K4me1 regions, suggesting that enhancers are particularly malleable during this developmental window. Furthermore, 86% (7,353 out of 8,528) of H3K4me1 regions change in accessibility during neuronal remodeling. Therefore, widespread temporal regulation of accessibility at enhancer regions is observed during neuronal remodeling.

Upon genome-wide examination of Shep-dependent differentially accessible regions, we found strong Shep-facilitated enhancer closing during neuronal remodeling. Overall, chromatin accessibility in Shep-depleted neurons shows substantially more changes in pupae (3,147 differential regions) than in larvae (331), consistent with pupal-specific transcriptome changes upon Shep depletion^11^. Considering only Shep-dependent accessibility changes in pupal neurons, we found 1,127 regions labeled by H3K4me1 that are normally kept inaccessible by Shep (Shep-inhibited). During normal neuronal development, these regions are accessible in larvae but become closed by the pupal stage (Fig. 4B and C). However, in Shep-depleted neurons, these regions gain additional accessibility in pupal compared to larval neurons, indicating greatly compromised closing of these putative enhancers. Intriguingly, these regions are most enriched for binding motifs for the zinc finger DNA-binding proteins, Klu transcription factor and GAGA-factor (GAF) (Fig. S4E and F), a protein that has been implicated in insulator activity^17,18^. While many enhancer regions failed to close during neuronal remodeling in the absence of Shep, we also identified 1,333 Shep-promoted regions labeled by H3K4me1 that are inaccessible in larval neurons but open during normal neuronal remodeling. Upon Shep depletion, these regions can only be partially opened, suggesting that Shep moderately contributes to the accessibility of these regions (Fig. 4B and D). For these regions, we obtained enrichment for motifs for Klu and the Sp1 family of transcription factors^19^ (Fig. S4F). By performing CUT&Tag of H3K27me3 with sorted control neurons from pupae, we observed more frequent Shep regulation of accessibility in either direction of H3K4me1-labeled enhancers (2,560) than H3K27me3-labeled inactive chromatin regions (1,305), despite similar genomic coverage by H3K4me1 (4.1e7 bp) and H3K27me3 (3.8e7 bp), indicating enhancer-biased Shep regulation of accessibility. Taken together, we conclude that Shep mainly mediates closing of numerous enhancers genome-wide during neuronal remodeling.

Finally, we verified that Shep-dependent changes in chromatin accessibility are functionally associated with gene expression changes genome-wide. *Myc*, another key downstream target that is repressed by Shep in pupal neurons^11,20^, displays temporally dynamic accessibility patterns at H3K4me1-enriched regions, some of which are also Shep-inhibited (Fig. S5A). Genome-wide, we found that Shep-inhibited accessibility changes are indeed statistically enriched for Shep-inhibited gene expression changes in pupal neurons (Fig. S5B). Statistically significant association between these two events is even more pronounced when restricted to H3K4me1-marked accessibility changes (Fig. 4E), suggesting that genome-wide enhancer closure is a key mechanistic step in transcriptional inhibition mediated by Shep. Among 82 genes inhibited by Shep for both expression and enhancer accessibility, Shep association is significantly enriched at enhancers of 26 genes (Fisher’s exact test (FET), *p=*1.7e-8, odds ratio=4.5), suggesting direct Shep inhibition of both enhancer accessibility and transcription. In contrast, correlation between Shep-promoted gene expression and Shep-regulated accessibility is either weaker (Fig. S5C and D) or absent (Fig. 4E; Fig. S5B) when enhancers are considered. These findings suggest that Shep inhibits chromatin accessibility, particularly enhancer accessibility, to temporally regulate gene expression during neuronal remodeling.

## DISCUSSION

Shep attenuates transcription of a critical developmental regulator during a specific window of neuronal remodeling by inhibiting E-P looping, which is itself dynamically controlled. Genome-wide, Shep reduces transcription of many downstream targets and limits chromatin accessibility of corresponding enhancer regions during neuronal remodeling. Although several transcription factors have been shown to inhibit E-P loop formation^6-8^, loss of looping in these cases is likely an indirect result of repressed enhancer and/or promoter activities. In these studies and our own work, it remains a challenge to distinguish between cause and consequence; however, we demonstrated that Shep does not alter the ability of *brat* enhancers or promoters to drive a reporter either *ex vivo* or *in vivo*. Thus, we propose that Shep is a dedicated anti-looping factor, which functions primarily by inhibiting E-P looping to regulate *brat* expression.

We speculate that Shep antagonism of E-P looping involves regulation of other DNA-binding proteins with respect to their chromatin association, as evidenced by changes of chromatin accessibility at enhancers. Our motif analysis of Shep-regulated enhancers identified Klu and the Sp1 family of transcription factors, two classes of zinc-finger proteins that are both enriched in the nervous system during metamorphosis^14^ when neuronal remodeling occurs. We also identified enrichment of a GA-rich motif known to be bound by the insulator-associated factor GAF, and this motif is present in the region 4 enhancer. Although GAF is an attractive candidate considering that it binds the *brat-F* promoter as well as enhancer regions 1 and 4 in chromatin of BG3 cells (Fig. S4G), the GA-rich motifs identified in our analyses are fairly generic repeated sequences that could alternatively be bound by other factors, such as CLAMP^21^ or AGO2^22^, for example. Future studies are required to interrogate the potential roles of these proteins in Shep mechanistic function. In addition, Shep is known to bind transcripts of its chromatin target genes^11,23^, raising the possibility that Shep may load onto chromatin during transcription and concomitantly regulate chromatin accessibility and E-P looping. Finally, specific coding or non-coding RNAs could be required for Shep function in this context, as RNA-binding is required for Shep antagonism of *gypsy* insulator activity^9^. Our results do not rule out the possibility that Shep also downregulates *brat* or other genes by posttranscriptional mechanisms. Given widespread evidence that E-P looping and enhancer accessibility are both frequently associated with gene activation, we predict that regulation of chromatin looping and enhancer accessibility are commonly utilized cellular mechanisms to control developmental gene expression programs.

Notably, our results provide the first evidence of temporal regulation of 3D chromatin organization that facilitates post-mitotic neuronal remodeling. Previous studies in *Drosophila* have identified a variety of genetic and cellular processes underlying neuronal remodeling^12,20,24^, and recent studies have begun to elucidate transcriptome dynamics throughout this developmental program^11,25^. However, fundamental mechanisms regulating 3D chromatin structure that result in temporal changes in gene expression during this essential process in any model system remain undefined. Our data reveal highly dynamic enhancer accessibility during normal neuronal maturation, similar to what has been observed during developmental differentiation in the mouse cerebellum^26^, suggesting conserved regulation of enhancer accessibility during neuronal maturation. We found that Shep inhibits enhancer accessibility during the temporal progression of neuronal remodeling, which corresponds to Shep chromatin association, Shep repression of chromatin looping, and temporal inhibition of the key target *brat* gene. Moderate expression changes of all *brat* isoforms as well as *brat* enhancer accessibility may result from changes occurring only in specific neurons within the total isolated population and/or transient changes in chromatin accessibility. Since we and others have observed that E-P looping is not necessarily sufficient to activate transcription^27^, it is likely that additional chromatin-related events must occur in order to achieve transcriptional activation. Widespread and conserved dynamics of enhancer accessibility is observed in the nervous system across organisms^26,28,29^; likewise, regulated E-P looping may also be utilized genome-wide during neuronal remodeling to regulate temporal gene expression. Finally, our findings suggest potential mechanisms underlying functions of human Shep orthologs, of which mutation and misexpression are associated with various neurological diseases, including ALS, Alzheimer’s, Parkinson’s, and schizophrenia^30^.

## Materials and Methods

### Fly strains

Fly stocks and crosses were grown on standard cornmeal-yeast-agarose medium at 25°C. Only female animals were used for experiments. Fly strains used include *elav-Gal4* (FBst0000458), *UAS-Dcr-2* (FBst0024650), and *UAS-shep-RNAi* (FBst0462204). For generation of the *phsp70-*GFP strains, region 1 or 4 was cloned into NotI restriction sites in the pEGFP-attB plasmid (*Drosophila* Genomics Resource Center). The resulting plasmids were integrated by *phiC31* integrase at the *attP40* locus by BestGene. To generate flies deleted for region 4 using CRISPR/Cas9 editing, two gRNAs CACTGTGCCAGAAAGTTCCA and AATGCACTGATTAACAGTAA were inserted into the BbsI sites of the plasmid pUC57 (a gift from B. Oliver, NIDDK) for gRNA expression after embryo injection. To delete endogenous region 4, genomic sequences flanking region 4 were inserted on either side of dsRed coding sequence into the plasmid pUC19 (a gift from B. Oliver, NIDDK) as a homologous repair template. Both plasmids were constructed by GenScript and were injected into *yw;;nos-Cas9(III-attP2)* flies by BestGene. Full plasmid sequences and primers used to validate the insertion location by sequencing are included in Table S1.

### Cell culture and transfection

BG3-c2 cells and S2 cells (*Drosophila* Genomics Resource Center) were cultured at 25°C in Schneider’s medium with 10% FBS or M3+BPYE medium with 10% FBS, respectively. BG3 medium was further supplemented with 10 µg/mL insulin. The MEGAscript T7 Kit (Thermo Fisher Scientific) was used to generate dsRNA, which was purified using NucAway Spin Columns (Thermo Fisher Scientific). The Amaxa Cell Line Nucleofector Kit V (Lonza) was used according to the manufacturer’s protocol to transfect constructs and dsRNA into 5-10 million BG3 or S2 cells using programs T30 or G30, respectively. Two µg of dsRNA targeting Shep or GFP was used to deplete Shep or serve as a control. One µg of each luciferase construct and 1 µg of Renilla control construct were co-transfected into cells to perform luciferase assays. Experiments were performed 4 d after transfection.

### Cloning of luciferase constructs

Luciferase cDNA was inserted between the XhoI and SpeI restriction sites of the pJET1.2 cloning vector. The 2 kb region upstream of *brat-A/E, brat-B/C,* or *brat*-*F* was amplified from Oregon-R genomic DNA and then inserted between the XhoI and EcoRI restriction sites upstream of the luciferase-encoding sequences. Candidate enhancer regions based on 4C-seq analysis were individually cloned and inserted upstream of the *brat-A/E, bratB/C*, or *brat-F* promoter at NotI restriction sites. Cloning primer sequences are listed in Table S1.

### Luciferase Assays

One and a half mL of cells from each transfection were spun at 600 x*g* for 10 min. Cell pellets were frozen at -80°C until analysis. Cells were resuspended in 250 µL nuclease-free water, and 75 µL were plated in triplicate in opaque 96-well plates. Then 75 µL of Dual-Glo Reagent (Promega) was added to each well, and the plate was incubated for 10 min at RT. Firefly luminescence was measured using a Spectramax II Gemini EM plate reader (Molecular Devices). Next, 75 µL of Dual-Glo Stop & Glo Reagent (Promega) was added to each well and incubated for 10 min at RT before measuring Renilla luminescence. Two to three biological replicates were performed per experiment.

### Chromatin conformation capture (3C)

For each replicate, 20 brains were dissected in Schneider’s medium containing 10% FCS and 10 µg/mL insulin. Formaldehyde was added to a final concentration of 2% and brains were fixed for 15 min at RT, then quenched with 0.125 M glycine for 10 min, followed by two 10 min rinses with washing buffer (50 mM Tris, 10 mM EDTA, 0.5 mM EGTA, 0.25% Triton-X100). Fixed brains were stored in storage buffer (10 mM Tris-HCl pH=8, 1 mM EDTA, 0.5 mM EGTA) at -80°C until all samples were pestle-homogenized in lysis buffer [10 mM NaCl, 0.2% NP-40, 10 mM Tris pH 8, and Mini Complete tablet (Roche)] and incubated at 37°C for 20 min. Nuclei were pelleted at 6,000 x*g* for 5 min, and the incubation was repeated once more. Nuclei were washed with digestion buffer (ThermoFisher Scientific ER0932 plus 0.2% NP-40), pelleted at 6,000 x*g* for 5 min and incubated with digestion buffer (ThermoFisher Scientific ER0932 plus 0.2% NP-40, 0.1% SDS) at 65°C for 30 min. Triton X-100 was added to a final concentration of 1%, and samples were further incubated at 37°C for 15 min. Ten percent of each sample was saved as an undigested control. Next, 200 U of BsRGI (ThermoFisher Scientific ER0932) was added to a final volume of 100 µL, and digestion was incubated for 2 d at 37°C. The restriction enzyme was inactivated at 65°C for 20 min, and another 10% of the sample was saved as a digested control. The digested sample was diluted with 400 µL T4 ligation buffer (NEB, M0202, 1% Triton X-100) and incubated at 37°C for 30 min, followed by an overnight incubation at 16°C with 3 µL T4 DNA ligase (NEB, M0202). Samples were purified with phenol-chloroform and used as 3C templates for Taqman-qPCR. The BAC CH321-86O1 (Chori) was used to generate 3C templates to normalize for primer efficiency, and primers targeting the *drl* locus were used to equalize loading across samples. Student’s *t* test was performed at \**p<*0.05. Sequences of the MGB (minor groove binder) Taqman probe, 3C primers, and loading control primers are included in Table S1.

### 4C-seq libraries

Generation of 4C-seq libraries in BG3 cells was performed using the brain 3C protocol except that BG3 cells were fixed with 1% formaldehyde for 10 min at RT, and the primary digestion enzyme used was Csp6I (ThermoFisher Scientific ER0211). Next, 3C templates were further processed according to a published 4C protocol^31^ with DpnII (NEB R0543) as the secondary digestion enzyme. Libraries were sequenced at the NIDDK Genomics Core Facility on an Illumina HiSeq 2500. Primers used to amplify 4C-seq libraries are documented in Table S1.

### 4C-seq analyses

Reads from the 4C assay were aligned to the dm6 genome using bowtie2 v2.3.5 with default parameters. Subsequent sam files were sorted and converted into bam files using the Samtools v1.9 view and sort commands, respectively. Sorted bam files were then used to perform the 4C-seq analysis with the FourCSeq v1.18.0 software package. Next, 4C-seq peak identification was performed by creating two dataframes containing the following metadata information: restriction enzymes, sequencing primers, reference genome ID, replicate information, viewpoint location and sorted bam file names. This metadata was used to create the *in silico* digested reference genome, extract the location of the viewpoint, and map its reads to both the reference genome and the fragment reference. Mapped reads were then counted using the ‘countFragmentOverlaps’ command in a strand specific manner. Because the restriction enzymes had already been trimmed, the *trim* parameter was set to 0, and the minimum-mapping quality was set to 20. Counts from both left and right fragment ends were combined using the ‘combineFragEnds’ command. Spikes and PCR artifacts were removed using ‘smoothCounts’, and z-scores were calculated for potential peaks along the default distance from the viewpoint using ‘getZScores’. Lastly, differential interacting fragments were identified using the ‘addPeaks’ function if at least one replicate had an adjusted *p* value of 0.01 and both replicates had a z-score larger than 3.

### FACS and ATAC-seq libraries

A published FACS procedure was used to dissociate and select GFP positive cells^11^. Cells were sorted on a FACSAria II machine at the Flow Cytometry Core of the National Heart, Lung and Blood Institute. Omni-ATAC-seq libraries were generated according to a detailed protocol^16^ with minor adjustments. Specifically, 2 × 10^5^ sorted GFP-positive neurons were used for each of three biological replicates, and DNA was phenol-chloroform extracted. Eleven total PCR cycles were performed to amplify libraries, which were subsequently double size-selected with AMPure XP beads (first with 0.6x volume, then with 1.2x volume) (Beckman Coulter). Finally, 50 bp paired-end sequencing was performed at the NIDDK Genomics Core Facility on an Illumina NextSeq 550 with the High Output mode.

### ATAC-seq computational analyses

Adapter sequences were trimmed from reads with cutadapt (v2.3; -a CTGTCTCTTATACACATCTCCGAGCCCACGAGAC -A CTGTCTCTTATACACATCTGACGCTGCCGACGA --minimum-length 18) and aligned to Flybase release dm6 reference with bowtie2 (v2.3.5; --very-sensitive, pair-end mode)^32^. Reads were depleted for those aligned to mitochondria (egrep -v chrM) and for multi-mapped reads with samtools (v1.9)^33^. Uniquely mapped reads were further depleted for PCR duplicates with picard (MarkDuplicates REMOVE_DUPLICATES=true) and computationally size-selected for inserts <150 bp to exclude nucleosome-related reads. ATAC-seq peak calling was performed with MACS2 (v2.2.6; pair-end mode -f BAMPE)^34^, and differential accessibility was called with the R package DiffBind v2.6.6 (edgeR, FDR<0.1 and lfc>0.3). Motif enrichment analyses were performed with AME 5.3.3^35^ and STREME 5.3.3^36^, and only motifs identified by both algorithms with known binding factors were reported.

### CUT&Tag libraries and analyses

After FACS sorting, collected neurons were used directly to generate CUT&Tag libraries following a protocol established by the Henikoff group (https://dx.doi.org/10.17504/protocols.io.bcuhiwt6) with minor changes. Specifically, 7.5 × 10^4^ sorted GFP-positive neurons were used for each of three biological replicates without crosslinking. The Tn5 was purchased from Epicypher (15-1017) and used at 1:20. The primary antibodies used were H3K27me3 (Cell Signaling Technology 9733), H3K4me1 (Abcam ab8895), and H3K27ac (Abcam ab4729). Sixteen total PCR cycles were performed to amplify libraries, which were re-purified (using 1.1x volume of AMPure XP beads) after pooling to remove primer dimers. Then 50 bp paired-end sequencing was completed at the NIDDK Genomics Core Facility on a NextSeq 550 using high-output. Computational analyses of CUT&Tag data were performed with the same pipeline used for ATAC-seq except that there was no computational size-selection of sequencing reads, and peaks were called by MACS2 (v2.2.6) in broad mode.

### Immunostaining and imaging

Immunostaining was performed as previously described^11^. The GFP primary antibody (ThermoFisher A10262) was used at 1:8,000, and guinea pig anti-Shep^11^ was used at 1:1,000. Images were taken as maximum-intensity z-series projections with a Zeiss780 confocal microscope.

### Statistics

All experiments were performed using three biological replicates unless noted. The *p* values of Fisher’s exact tests were calculated with R v3.6.1 and reported as two-tailed values at \*\*\**p*<0.001, \*\**p*<0.01 and \**p*<0.05. Averaged values are reported as mean±sem unless otherwise noted.

## Acknowledgements

We thank S. Wang for assistance with custom MA plots; L. Benner for guidance on CRISPR; and A. Dean, G. Blobel, and members of the Lei laboratory for comments on the manuscript.

## Author contributions

Conceptualization: D.C., C.E.M., E.P.L.; Data Curation: D.C., C.E.M., B.R.; Formal Analysis:

D.C., C.E.M., B.R.; Funding acquisition: E.P.L., D.C.; Investigation: D.C., C.E.M.; Methodology:

D.C., C.E.M., B.R.; Project administration: D.C., E.P.L.; Resources: E.P.L.; Software: D.C., B.R.;

Supervision: E.P.L.; Validation: D.C., C.E.M., E.P.L.; Visualization: D.C., C.E.M., B.R.; Writing – original draft: D.C.; Writing – review & editing: D.C., C.E.M., B.R., E.P.L.

## Data and materials availability

The 4C-seq, ATAC-seq, CUT&Tag, and ChIP-seq data have been deposited in the Gene Expression Omnibus database under accession number GSE154645; All data are available in the main text or supplementary materials.

## References

1 Ghavi-Helm, Y. et al. Enhancer loops appear stable during development and are associated with paused polymerase. Nature 512, 96–100, doi:10.1038/nature13417 (2014).

2 Noordermeer, D. et al. The dynamic architecture of Hox gene clusters. Science 334, 222–225, doi:10.1126/science.1207194 (2011).

3 Deng, W. et al. Controlling long-range genomic interactions at a native locus by targeted tethering of a looping factor. Cell 149, 1233–1244, doi:10.1016/j.cell.2012.03.051 (2012).

4 Kim, J. H. et al. LADL: light-activated dynamic looping for endogenous gene expression control. Nat Methods 16, 633–639, doi:10.1038/s41592-019-0436-5 (2019).

5 Deng, W. et al. Reactivation of developmentally silenced globin genes by forced chromatin looping. Cell 158, 849–860, doi:10.1016/j.cell.2014.05.050 (2014).

6 Chopra, V. S., Kong, N. & Levine, M. Transcriptional repression via antilooping in the Drosophila embryo. Proc Natl Acad Sci U S A 109, 9460–9464, doi:10.1073/pnas.1102625108 (2012).

7 Kim, Y. H. et al. Rev-erbalpha dynamically modulates chromatin looping to control circadian gene transcription. Science 359, 1274–1277, doi:10.1126/science.aao6891 (2018).

8 McClellan, M. J. et al. Modulation of enhancer looping and differential gene targeting by Epstein-Barr virus transcription factors directs cellular reprogramming. PLoS Pathog 9, e1003636, doi:10.1371/journal.ppat.1003636 (2013).

9 Chen, D., Brovkina, M., Matzat, L. H. & Lei, E. P. Shep RNA-Binding Capacity Is Required for Antagonism of gypsy Chromatin Insulator Activity. G3 (Bethesda) 9, 749–754, doi:10.1534/g3.118.200923 (2019).

10 Matzat, L. H., Dale, R. K., Moshkovich, N. & Lei, E. P. Tissue-specific regulation of chromatin insulator function. PLoS Genet 8, e1003069, doi:10.1371/journal.pgen.1003069 (2012).

11 Chen, D., Dale, R. K. & Lei, E. P. Shep regulates Drosophila neuronal remodeling by controlling transcription of its chromatin targets. Development 145, doi:10.1242/dev.154047 (2018).

12 Chen, D., Qu, C., Bjorum, S. M., Beckingham, K. M. & Hewes, R. S. Neuronal remodeling during metamorphosis is regulated by the alan shepard (shep) gene in Drosophila melanogaster. Genetics 197, 1267–1283, doi:10.1534/genetics.114.166181 (2014).

13 Olesnicky, E. C., Bhogal, B. & Gavis, E. R. Combinatorial use of translational co-factors for cell type-specific regulation during neuronal morphogenesis in Drosophila. Dev Biol 365, 208–218, doi:10.1016/j.ydbio.2012.02.028 (2012).

14 Celniker, S. E. et al. Unlocking the secrets of the genome. Nature 459, 927–930, doi:10.1038/459927a (2009).

15 Yanez-Cuna, J. O. et al. Dissection of thousands of cell type-specific enhancers identifies dinucleotide repeat motifs as general enhancer features. Genome Res 24, 1147–1156, doi:10.1101/gr.169243.113 (2014).

16 Corces, M. R. et al. An improved ATAC-seq protocol reduces background and enables interrogation of frozen tissues. Nat Methods 14, 959–962, doi:10.1038/nmeth.4396 (2017).

17 Ohtsuki, S. & Levine, M. GAGA mediates the enhancer blocking activity of the eve promoter in the Drosophila embryo. Genes Dev 12, 3325–3330, doi:10.1101/gad.12.21.3325 (1998).

18 Schweinsberg, S. et al. The enhancer-blocking activity of the Fab-7 boundary from the Drosophila bithorax complex requires GAGA-factor-binding sites. Genetics 168, 1371–1384, doi:10.1534/genetics.104.029561 (2004).

19 Kaczynski, J., Cook, T. & Urrutia, R. Sp1-and Kruppel-like transcription factors. Genome Biol 4, 206, doi:10.1186/gb-2003-4-2-206 (2003).

20 Chen, D., Gu, T., Pham, T. N., Zachary, M. J. & Hewes, R. S. Regulatory Mechanisms of Metamorphic Neuronal Remodeling Revealed Through a Genome-Wide Modifier Screen in Drosophila melanogaster. Genetics 206, 1429–1443, doi:10.1534/genetics.117.200378 (2017).

21 Soruco, M. M. et al. The CLAMP protein links the MSL complex to the X chromosome during Drosophila dosage compensation. Genes Dev 27, 1551–1556, doi:10.1101/gad.214585.113 (2013).

22 Moshkovich, N. et al. RNAi-independent role for Argonaute2 in CTCF/CP190 chromatin insulator function. Genes Dev 25, 1686–1701, doi:10.1101/gad.16651211 (2011).

23 Dale, R. K., Matzat, L. H. & Lei, E. P. metaseq: a Python package for integrative genome-wide analysis reveals relationships between chromatin insulators and associated nuclear mRNA. Nucleic Acids Res 42, 9158–9170, doi:10.1093/nar/gku644 (2014).

24 Yaniv, S. P. & Schuldiner, O. A fly’s view of neuronal remodeling. Wiley Interdiscip Rev Dev Biol 5, 618–635, doi:10.1002/wdev.241 (2016).

25 Alyagor, I. et al. Combining Developmental and Perturbation-Seq Uncovers Transcriptional Modules Orchestrating Neuronal Remodeling. Dev Cell 47, 38–52 e36, doi:10.1016/j.devcel.2018.09.013 (2018).

26 Frank, C. L. et al. Regulation of chromatin accessibility and Zic binding at enhancers in the developing cerebellum. Nat Neurosci 18, 647–656, doi:10.1038/nn.3995 (2015).

27 Ghavi-Helm, Y. et al. Highly rearranged chromosomes reveal uncoupling between genome topology and gene expression. Nat Genet 51, 1272–1282, doi:10.1038/s41588-019-0462-3 (2019).

28 Reddington, J. P. et al. Lineage-Resolved Enhancer and Promoter Usage during a Time Course of Embryogenesis. Dev Cell 55, 648–664 e649, doi:10.1016/j.devcel.2020.10.009 (2020).

29 Nord, A. S. & West, A. E. Neurobiological functions of transcriptional enhancers. Nat Neurosci 23, 5–14, doi:10.1038/s41593-019-0538-5 (2020).

30 Schachtner, L. T. et al. Drosophila Shep and C. elegans SUP-26 are RNA-binding proteins that play diverse roles in nervous system development. Dev Genes Evol 225, 319–330, doi:10.1007/s00427-015-0514-3 (2015).

31 Nazer, E., Dale, R. K., Chinen, M., Radmanesh, B. & Lei, E. P. Argonaute2 and LaminB modulate gene expression by controlling chromatin topology. PLoS Genet 14, e1007276, doi:10.1371/journal.pgen.1007276 (2018).

32 Langmead, B. & Salzberg, S. L. Fast gapped-read alignment with Bowtie 2. Nat Methods 9, 357–359, doi:10.1038/nmeth.1923 (2012).

33 Li, H. et al. The Sequence Alignment/Map format and SAMtools. Bioinformatics 25, 2078–2079, doi:10.1093/bioinformatics/btp352 (2009).

34 Zhang, Y. et al. Model-based analysis of ChIP-Seq (MACS). Genome Biol 9, R137, doi:10.1186/gb-2008-9-9-r137 (2008).

35 McLeay, R. C. & Bailey, T. L. Motif Enrichment Analysis: a unified framework and an evaluation on ChIP data. BMC Bioinformatics 11, 165, doi:10.1186/1471-2105-11-165 (2010).

36 Bailey, T. L. STREME: Accurate and versatile sequence motif discovery. Bioinformatics, doi:10.1093/bioinformatics/btab203 (2021).

