## Supplemental figures 1-5 for "Temporal inhibition of chromatin looping and enhancer accessibility during neuronal remodeling"

### Fig. S1

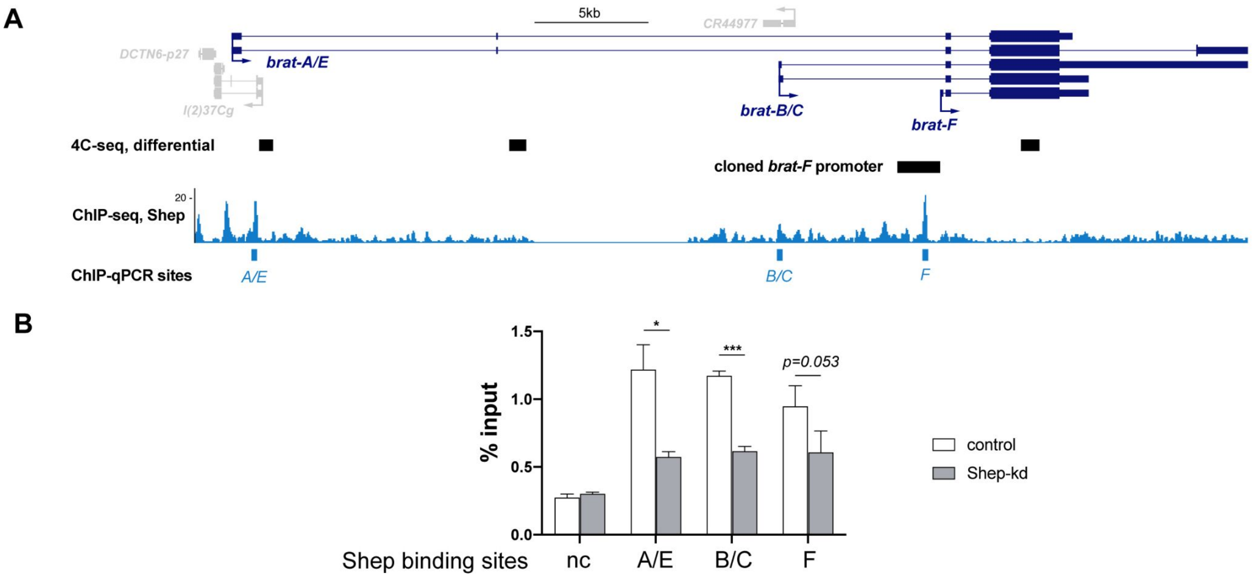


**Fig. S1. Validation of Shep chromatin association in BG3 cells.** BG3 cells treated with GFP or Shep dsRNA were used for ChIP-qPCR quantification of Shep binding near three *brat* isoform promoters. Significant or marginally significant reduction of Shep binding was observed for all three loci upon Shep depletion. **A)** Screenshot and **B)** directed ChIP-qPCR quantification of three biological replicates.

**Fig. S2.**

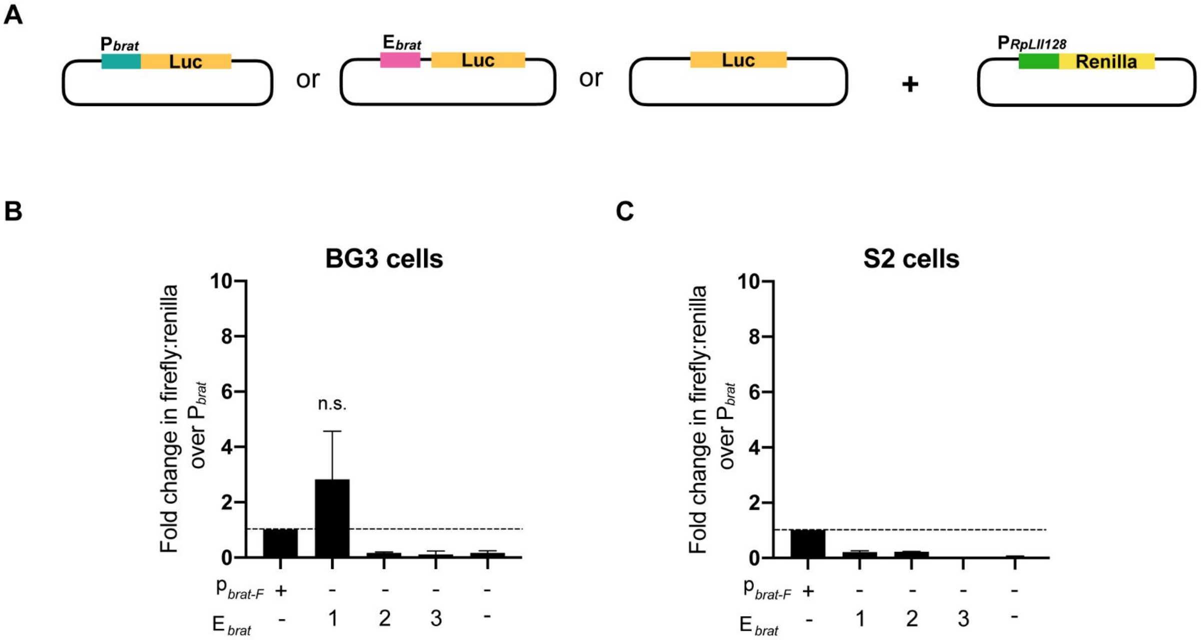


**Fig. S2. Shep does not repress activity of the region 1 enhancer or any *brat* promoters. (A)** Control constructs with luciferase cloned downstream of the *brat-F* promoter or individual enhancer candidates alone were co-transfected with the Renilla control construct. **(B)** Fold change over *brat-F* promoter alone driven expression of Renilla-normalized luciferase activity in BG3 cells. None of the enhancer candidates alone drive stronger luciferase than the *brat-F* promoter in BG3 cells. **(C)** Identical transfections as in (B) in S2 cells.

**Fig. S3.**

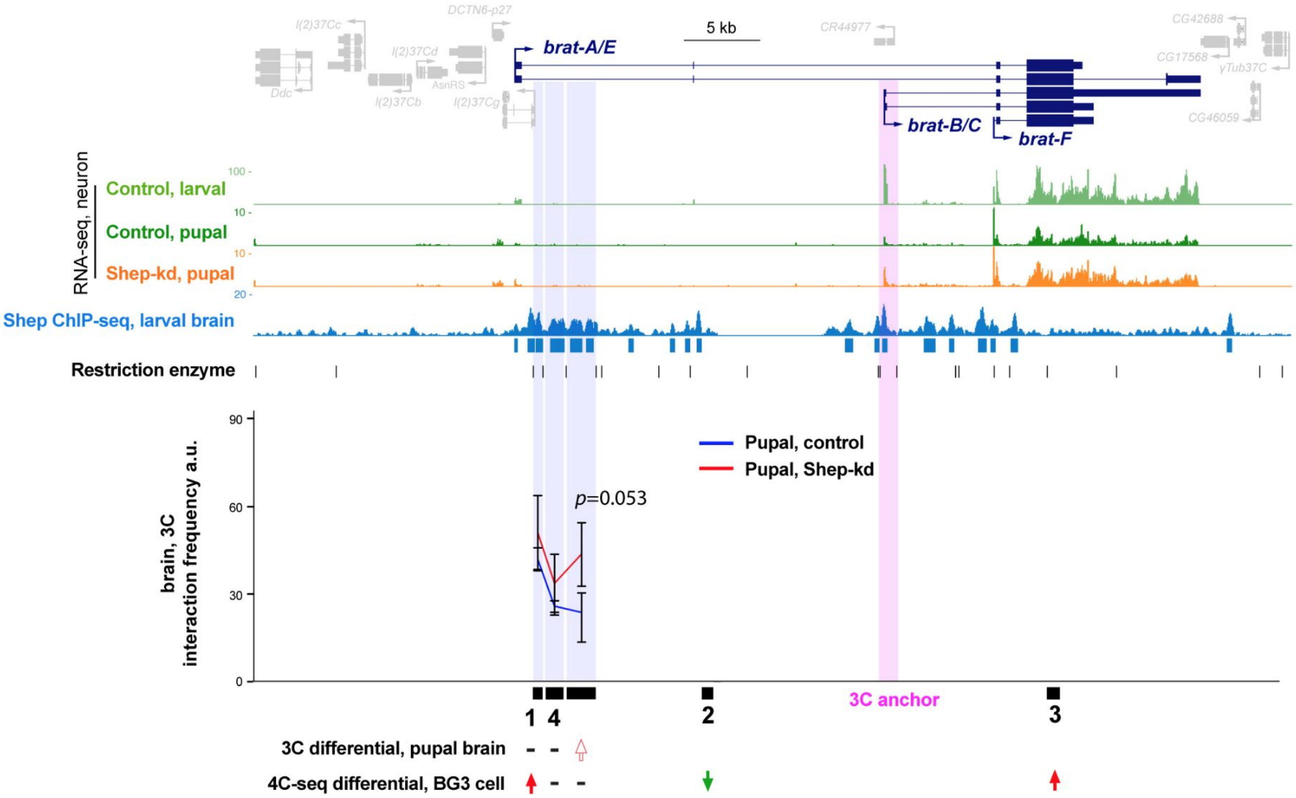


**Fig. S3.** Pupal brain 3C assays with the *brat-B/C* promoter as the anchor. Three biological replicates were included for each genotype. Enhancer 1 and enhancer 4 do not show changed interaction frequencies with the *brat-B/C* promoter. The 3' region shows a marginally significant increase of interaction frequency. Note difference in 3C scale compared to Figure 3.

Fig. S4.

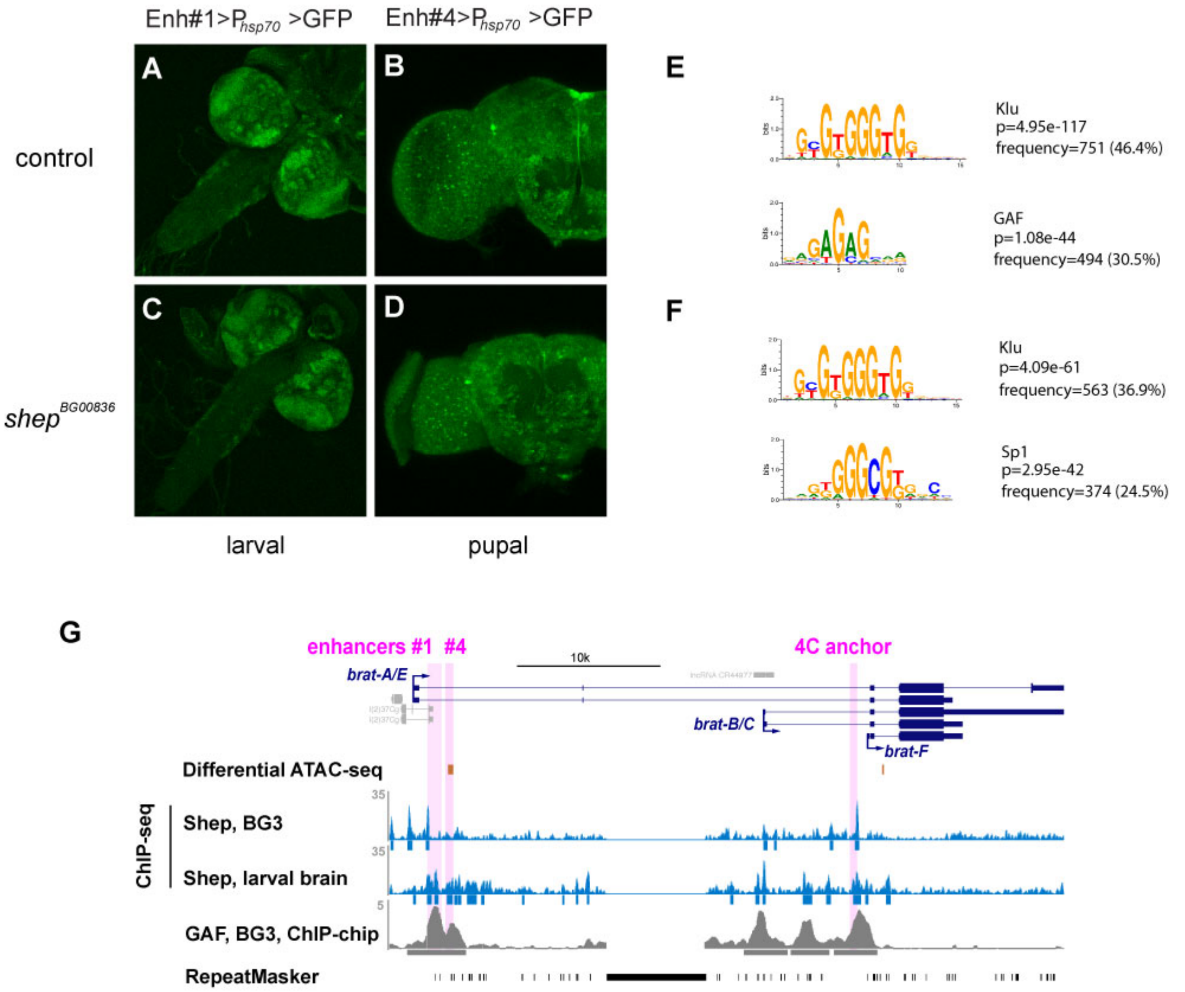


**Fig. S4. Region 1 and 4 enhancer activities and accessibility are unchanged in Shep-depleted pupal brains and neurons. (A-D)** GFP expression driven by region 1 or 4 directly upstream of the *hsp70* minimal promoter in larval or pupal brains, respectively, in control or strong loss-of-function *shep* mutant brains. **(E)** Top motifs identified by motif enrichment analyses using algorithms AME and STREME of H3K4me1-labeled Shep-inhibited accessible regions in pupal neurons. Frequency indicates the proportion of Shep-inhibited enhancers that harbor respective motifs. **(F)** Motifs identified for enhancers of Shep-promoted accessibility. **(G)** GAF associates with regulatory elements and promoters of *brat* in BG3 cells.

**Fig. S5.**

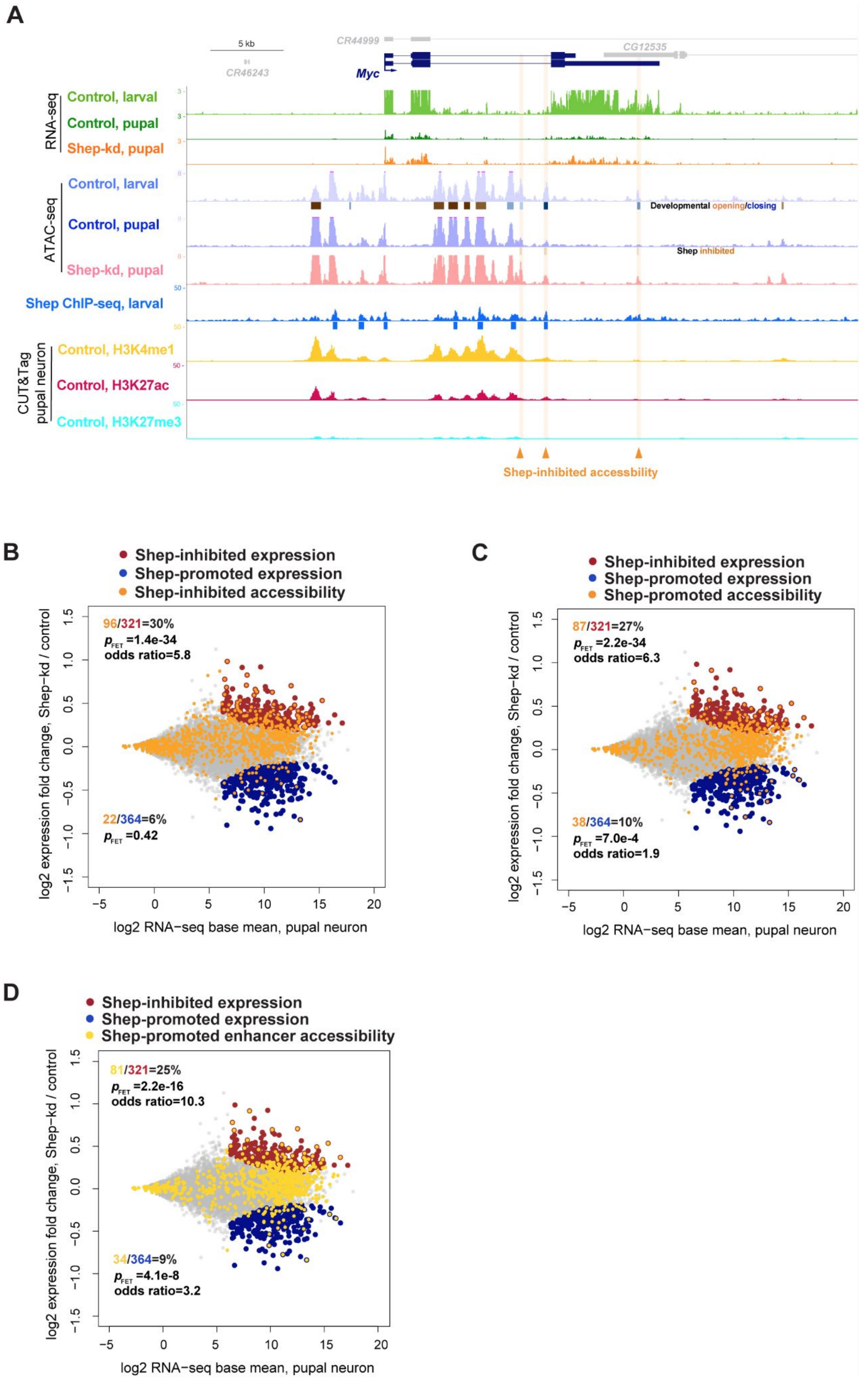
A
5 kb
CR44999
CR46243
CG12535
Myc
RNA-seq
Control, larval
Control, pupal
Shep-kd, pupal
ATAC-seq
Control, larval
Developmental opening/closing
Control, pupal
Shep inhibited
Shep-kd, pupal
Shep ChIP-seq, larval
CUT&Tag pupal neuron
Control, H3K4me1
Control, H3K27ac
Control, H3K27me3
Shep-inhibited accessbility
B
Shep-inhibited expression
Shep-promoted expression
Shep-inhibited accessibility
96/321=30%
pFET =1.4e-34
odds ratio=5.8
22/364=6%
pFET =0.42
log2 expression fold change, Shep−kd / control
log2 RNA−seq base mean, pupal neuron
C
Shep-inhibited expression
Shep-promoted expression
Shep-promoted accessibility
87/321=27%
pFET =2.2e-34
odds ratio=6.3
38/364=10%
pFET =7.0e-4
odds ratio=1.9
log2 expression fold change, Shep−kd / control
log2 RNA−seq base mean, pupal neuron
D
Shep-inhibited expression
Shep-promoted expression
Shep-promoted enhancer accessibility
81/321=25%
pFET =2.2e-16
odds ratio=10.3
34/364=9%
pFET =4.1e-8
odds ratio=3.2
log2 expression fold change, Shep−kd / control
log2 RNA−seq base mean, pupal neuron

**Fig. S5. Genome-wide association between Shep-regulated expression and accessibility. (A)** Pupal-specific Shep inhibition of *Myc* expression (orange track, $p$=5.6e-3, fold change=1.5) and accessibility of H3K4me1-labeled regions (orange shading, FDR=0.06, 0.05, and 0.03, respectively; fold change=1.3, 1.3 and 1.5, respectively). **(B)** Overlap between Shep-dependent expression and Shep-inhibited accessibility in pupal neurons. Genes inhibited for expression by Shep are enriched for Shep-inhibited accessible regions (red and orange). **(C-D)** Overlap between Shep-dependent expression and Shep-promoted accessibility in pupal neurons.
